# Implicit sensorimotor adaptation is preserved in Parkinson’s Disease

**DOI:** 10.1101/2022.03.11.484047

**Authors:** Jonathan S. Tsay, Tara Najafi, Lauren Schuck, Tianhe Wang, Richard B. Ivry

**Author notes:** These authors contributed equally to this work (Jonathan S. Tsay, Tara Najafi). **Corresponding author Information:** Name: Jonathan Tsay, Address: 2121 Berkeley Way, Berkeley, CA 94704.

## Abstract

Our ability to enact successful goal-directed actions involves multiple learning processes. Among these processes, implicit motor adaptation ensures that the sensorimotor system remains finely tuned in response to changes in the body and environment. Whether Parkinson’s Disease (PD) impacts implicit motor adaptation remains a contentious area of research: whereas multiple reports show impaired performance in this population, many others show intact performance. While there are a range of methodological differences across studies, one critical issue is that performance in many of the studies may reflect a combination of implicit adaptation and strategic re-aiming. Here, we revisited this controversy using a visuomotor task designed to isolate implicit adaptation. In two experiments, we found that adaptation in response to a wide range of visual perturbations (3° - 45°) was similar in PD and matched control participants. Moreover, in a meta-analysis of previously published work, we found that the mean effect size contrasting PD and controls across 16 experiments was not significant. Together, these analyses indicate that implicit adaptation is preserved in PD, offering a fresh perspective on the role of the basal ganglia in sensorimotor learning.

**Significance statement:** Among multiple motor learning processes, implicit adaptation ensures that our motor system remains exquisitely calibrated. Whether Parkinson’s disease affects implicit motor adaptation has been a point of controversy. We revisited this issue using a visuomotor task designed to isolate implicit adaptation and found that individuals with PD and matched controls showed indistinguishable performance. A meta-analysis based on data from 16 previous experiments yielded a similar null result, strongly supporting the notion that implicit adaptation is preserved in PD.

## Introduction

Parkinson’s Disease, a neurodegenerative disorder primarily affecting the basal ganglia circuitry, is thought to impair the ability to acquire and adapt skilled movements (1–3). Evidence of this comes from research involving a wide range of motor learning tasks including sequence learning (4–8), sensorimotor adaptation (9,10), and skill acquisition (11). Coupled with evidence implicating the basal ganglia in habitual behavior (12,13), this body of work has motivated the idea that the basal ganglia is essential for the acquisition, refinement, and automatization of skilled movement (14)

In the present study, we focus on one form of sensorimotor learning, visuomotor adaptation. This type of learning keeps the motor system exquisitely calibrated by automatically adjusting the sensorimotor map in response to errors between the expected and actual sensory feedback (15,16). A number of studies have shown that individuals with PD are impaired on visuomotor adaptation tasks (17–19); specifically, when a perturbation is imposed on the visual feedback, participants with PD fail to show the same level of compensation (i.e., movement in the opposite direction of the perturbation) as control participants. However, recent studies have revealed the operation of multiple learning processes even in seemingly simple tasks such as visuomotor adaptation. In particular, implicit adaptation may be supplemented by the use of volitional aiming. For example, to nullify an angular perturbation of the visual feedback, participants may aim away from the target (20,21). As such, the impaired performance of PD participants on adaptation tasks may not reflect disruption to implicit adaptation per se, but an impairment in other learning processes such as strategic aiming (see also, (22)).

To address this concern, we assessed PD and matched controls in a visuomotor adaptation task that isolates implicit motor adaptation (23). In this task, participants reach to a visual target and receive cursor feedback that follows a trajectory defined relative to the target and, importantly, is not contingent on the position/trajectory of the participant’s actual movement. Participants are fully informed of this manipulation and instructed to always reach directly to the target while ignoring the visual feedback. Despite these instructions, the mismatch introduced between the position of the target and the visual cursor induces an automatic adaptive response, causing a drift in movement direction away from the target and in the opposite direction of the cursor. These motor corrections are not the result of re-aiming; indeed, participants are oblivious to their change in their behavior (24,25).

We tested individuals with PD in two web-based experiments with this form of non-contingent feedback (for a validation of our web-based platform, see (26,27)). We complemented our empirical work with a meta-analysis, reviewing the results from 16 visuomotor adaptation experiments that allowed for a comparison between PD and control groups on a measure of adaptation that is unlikely to be significantly contaminated by other learning processes. Taken together, we sought to provide a thorough re-examination of the impact of PD on implicit motor adaptation.

## Methods

### Ethics Statement

All participants gave written informed consent in accordance with policies approved by UC Berkeley’s Institutional Review Board. Participation in the study was conducted online and in exchange for monetary compensation.

### Participants

Individuals diagnosed with Parkinson’s Disease (PD, total N = 25) were recruited via our clinical database (Table S1). The database is composed of individuals from around the country who have responded to online advertisements distributed by leaders of local support groups. We follow-up with a video call to describe the project in detail. For those who wish to participate, we obtain a medical history, neurological evaluation of PD symptoms using the motor component of the Unified Parkinson’s Disease Rating Scale (28), and evaluation of general cognitive status with the Montreal Cognitive Assessment (29,30). The UPDRS and MoCA were modified for online administration (27). For the UPDRS, we eliminated one item (postural stability) and three items were scored based on self-reports (arising from chair, posture, gait) due to concerns about safety during remote examination. For the MoCA, we eliminated the Alternating Trail Making item since it requires a paper copy. As shown in Table 1, the PD participants tended to have mild (0 - 20) to moderate (21–40) motor impairments. Four of the PD participants exhibited mild cognitive impairment (MoCA score < 26).

**Table 1:** PD and matched control participants. Participants were either right-handed (R), left-handed (L), or ambidextrous (A). With our modified test, UPDRS scores can range from 0 to 40 (where high numbers indicate greatest symptomology) and MoCA scores can range from 0 to 30 (where low numbers indicate greatest impairment). Mean (SD) is provided.

| Exp | Group | N | Age | Gender | Handedness | Years of Education | MoCA | UPDRS |
| --- | --- | --- | --- | --- | --- | --- | --- | --- |
| 1 | PD | 18 | 65.7 (7.6) | 8 M, 10 F | 14R, 3L, 1A | 17.4 (3.3) | 27.8 (1.8) | 16.6 (5.1) |
|  | Control | 18 | 64.1 (6.9) | 8 M, 10 F | 14R, 3L, 1A | 15.3 (2.2) | -- | -- |
| 2 | PD | 16 | 66.9 (7.2) | 7 M, 9 F | 13R, 3L | 17.6 (3.2) | 27.4 (1.9) | 16.7 (5.0) |
|  | Control | 16 | 65.6 (7.2) | 7 M, 9 F | 13R, 3L | 15.4 (2.6) | -- | -- |

9 of the 25 PD participants were tested only in Experiment 1, 7 were tested only in Experiment 2, and 9 were tested in both Experiments 1 and 2. The two experiments were separated by at least six months. The PD participants were on their normal medication schedule at the time of the video call and online testing. 34 control participants were recruited via Prolific, a website for online participant recruitment. Drawing on our database, we recruited individuals to match the PD participants in terms of age, gender, and handedness (Table 1). Each control participant was tested in a single experiment. The PD group had more years of education than the control group (Exp 1: *t*_30_ = 2.1, *p* = 0.04, [0.06, 4.3], *D* = 0.7; Exp 2: *t*_34_ = 2.3, *p* = 0.02, [0.2,4.0], *D* = 0.8). We did not include years of education in our selection criteria given that implicit motor adaptation is unlikely to depend on this measure. Nonetheless, given the group difference in education, we examined the possible impact of years of education as a covariate of interest in both experiments.

### Apparatus and Procedure

Participants used their own laptop or desktop computer to access a customized webpage that hosted the experiment (26,31,32). Participants used their computer trackpad or mouse to perform the reaching task (sampling rate typically ~60 Hz). The size and position of stimuli were scaled based on each participant’s screen size. For ease of exposition, the stimuli parameters reported below are for a typical monitor size of 13’’ with screen resolution of 1366 x 768 pixels (33).

Reaching movements were performed by using the computer trackpad or mouse to move the cursor across the monitor. Each trial involved a planar movement from the center of the workspace to a visual target. The center position was indicated by a white circle and the target location was indicated by a blue circle (both 0.5 cm in diameter). On the typical monitor, the radial distance from the start location to the target was 6 cm. The target appeared at one of two locations on an invisible virtual circle (135° = upper left quadrant; 315° = lower right quadrant). The movement involved some combination of joint rotations about the arm, wrist, and/or finger depending on whether the trackpad or mouse was used. In our prior validation work using this online interface and procedure, the exact movement and the exact device used did not impact measures of performance or learning on visuomotor adaptation tasks (26).

To initiate each trial, the participant moved the cursor, represented by a white dot (0.5 cm in diameter) into the start location. Feedback during this initialization phase was only provided when the cursor was within 2 cm of the start circle. Once the participant maintained the cursor in the start position for 500 ms, the target appeared. The participant was instructed to reach, attempting to rapidly “slice” through the target. We did not impose any constraints on reaction time. However, to discourage mid-movement corrections, we required that the movement be completed within 500 ms. If the movement time exceeded this criterion, the message “Too Slow” was displayed in the center of the screen for 750 ms.

We assumed that the stimulus display (vertical) and movement (horizontal) were in orthogonal planes. Given this assumption, the feedback cursor during the center-out movement could take one of three forms: Congruent feedback, rotated non-contingent feedback, and no-feedback. During congruent feedback trials, the movement of the visual feedback was congruent with the direction of the hand (e.g., rightward movement of hand produced a rightward movement of cursor; forward movement of hand produced an upward movement of cursor). During rotated non-contingent feedback trials (Figure 1a; Figure 2a, 2d), the cursor moved at a specified angular offset relative to the position of the target, regardless of the movement direction of the hand. The radial position of the cursor corresponded to that of the hand up to 6 cm, at which point the cursor position was frozen for 50 ms before disappearing. During no-feedback trials, the feedback cursor was extinguished as soon as the hand left the start circle and remained off for the entire reach. During the return phase after the movement, the (congruent) cursor was provided when the participant moved within 2 cm of the start circle.

**Figure 1:**
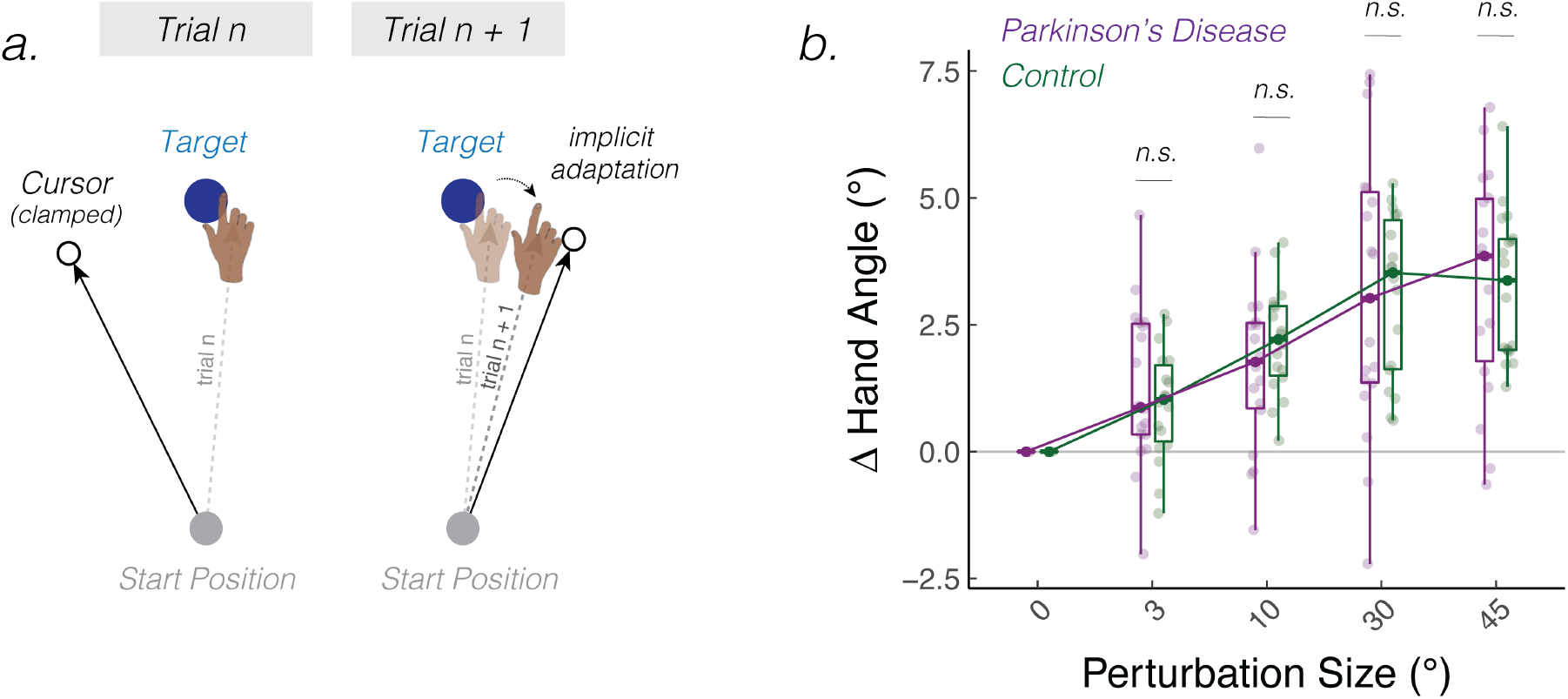
Parkinson’s Disease does not impact implicit adaptation in response to a wide range of error sizes. **(a)** Schematic of the task. The cursor feedback (hollow black circle) was rotated relative to the target, independent of the position of the participant’s hand. The size of the rotation was varied randomly on a trial-by-trial basis. **(b)** Average change in hand angle on the subsequent trial is plotted as a function of the rotation size for PD (dark magenta) and Control (green) participants. Box plots denote the median (thick horizontal lines), quartiles (1st and 3rd, the edges of the boxes), and extrema (min and max, vertical thin lines). The individual means are shown as translucent circles. *n.s.* denotes that the group comparison between PD and controls is not significant. Note that the 0° perturbation was not included in the actual experiment. It is plotted for illustrative purposes to highlight the slope of the motor correction function.

**Figure 2:**
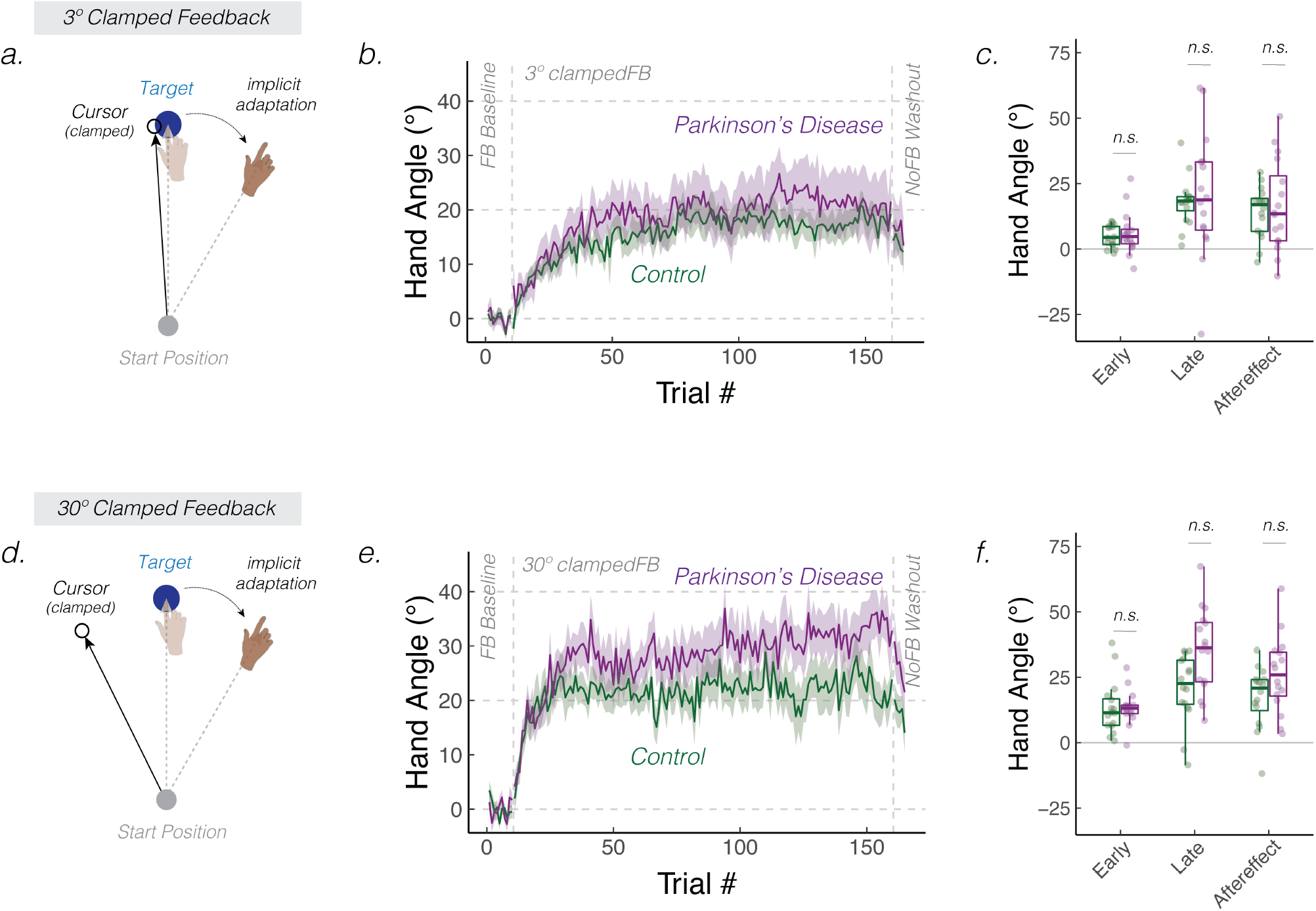
Parkinson’s Disease does not impact learning functions in response to clamped feedback. **(a, d)** Schematic of the clamped feedback task. The cursor feedback (hollow black circle) followed an invariant trajectory, rotated by either 3° **(a)** or 30° **(d)** relative to the target. The rotation size remained invariant over a block of 110 trials, with the order and sign of the rotation size counterbalanced across participants. Participants were instructed to always move directly to the target (blue circle) and ignore the visual feedback. The translucent and solid colors display hand position early and late in adaptation, respectively. **(b, e)** Mean time courses of hand angle for the 3° and 30° conditions for the PD (dark magenta) and Control (green) participants. Data for each participant were baseline subtracted relative to mean hand angle during the baseline phase with veridical feedback. Shaded region denotes SEM. **(c, f)** Average hand angles during early and late phases of the perturbation block, and during the no-feedback aftereffect block. Box plots denote the median (thick horizontal lines), quartiles (1st and 3rd, the edges of the boxes), and extrema (min and max, vertical thin lines). The mean for each participant is shown as translucent circles. *n.s.* denotes that the group comparison between PD and Controls is not significant

### Experiment 1: The Impact of PD on Implicit Adaptation

PD and Control participants (N = 18 per group) completed an adaptation task in which we examined the trial-by-trial response to non-contingent feedback (34,35). To familiarize participants with the apparatus and task requirements, the experiment began with a short baseline phase (10 trials) with congruent feedback. This was followed by 220 trials with non-contingent feedback in which the angular offset of the feedback cursor from the target varied from trial to trial, both in direction (clockwise - or counterclockwise +) and magnitude (3°, 10°, 30°, 45°). There were 26-28 trials per perturbation size provided in a random, zero-mean order (i.e., every perturbation was repeated at least once every 8 trials) to prevent any systematic drifts in hand angle too far away from the target.

Testing at the two target locations was varied across blocks, with one location used for trials 1-120 and the other location for trials 121-230 (i.e., each half contained 10 baseline congruent feedback trials and 110 non-contingent feedback trials). The target order was counterbalanced across participants. (We included the two locations to maintain a similar procedure in Experiments 1 and 2 – see *Experiment 2* methods below.)

Prior to the start of each baseline block, the instruction “Move directly to the target as fast and accurately as you can” appeared on the screen. Prior to the start of each perturbation block, the instructions were modified to read: “The white cursor will no longer be under your control. Please ignore the white cursor and continue to aim directly towards the target.” To clarify the invariant nature of the clamped feedback, three demonstration trials were provided before the first perturbation block. On all three trials, the target appeared straight ahead (90° position), and the participant was told to reach to the left (demo 1), to the right (demo 2), and backward (demo 3). On all three of these demonstration trials, the cursor moved in a straight line, 90° offset from the target. In this way, the participant could see that the spatial trajectory of the cursor was unrelated to their own reach direction.

Hand angle was defined as the position of the hand relative to the target when movement amplitude reached 6 cm. Due to data storage constraints, we opted not to record the entire movement trajectory since pilot data indicated that movement trajectories were relatively straight with minimal mid-movement corrections. As our key measure of trial-by-trial adaptation, we calculated the change in hand angle on trial n + 1 as a function of the clamped rotation size on trial n. The mean trial-by-trial change in hand angle was calculated for each perturbation size (combing both clockwise and counterclockwise directions). The mean data were submitted to a linear mixed effect model (R statistical package: lmer), with Rotation Size (3°, 10°, 30°, 45°) and Group (PD or Control) as fixed effects and Participant ID as a random effect. We also included Years of Education as a covariate, since this factor differed between groups. To visualize the data, the change in hand angle values with counterclockwise rotations were flipped, such that a positive hand angle corresponds to an angle in the opposite direction of the rotated feedback, the direction of movement expected from implicit adaptation.

### Experiment 2: The Impact of PD on the Upper Bound of Implicit Adaptation

As a second test of the effect of PD on implicit adaptation, we measured the cumulative effects of learning, an approach that should magnify any subtle differences between groups. To test this, we used a clamp of a fixed size and direction throughout the perturbation block, allowing learning to accumulate in one direction (i.e., the opposite direction of the perturbation). With this method, our focus was on the upper bound of adaptation and the aftereffect.

PD and Control participants (N = 16 per group) completed this task. To familiarize participants with the apparatus and task requirements, the experiment began with a short baseline block (10 trials) with congruent feedback. This was followed by a clamped non-contingent feedback perturbation block for 150 trials and a no-feedback aftereffect block for 5 trials. All three phases were then repeated at a different target location, amounting to a total of 330 trials

The target again appeared in one of two locations (135°, upper left quadrant; 315°, lower right quadrant). These two targets were spaced 180° apart to eliminate generalization of learning between target locations (23,36,37). One target was paired with a small (3°) perturbation and the other was paired with a large (30°) perturbation. The pairing between targets and clamp sizes, the direction of the clamped rotation (clockwise - or counterclockwise +), and their sequential order in the blocked design were counterbalanced across participants.

Task instructions were similar to that of Experiment 1: Prior to the baseline block, the participant was instructed to “Move directly to the target as fast and accurately as you can”. Prior to the non-contingent clamp block, the instructions were modified to read: “The white cursor will no longer be under your control. Please ignore the white cursor and continue to aim directly towards the target.” We again included three demonstration trials prior to the first clamped perturbation block, with the procedure identical to that of Experiment 1. Prior to the no-feedback aftereffect block, participants were again instructed to “Move directly to the target as fast and accurately as you can.”

To evaluate the magnitude of adaptation, the hand angle for each trial was calculated with respect to the participant’s idiosyncratic baseline bias (38,39). Baseline was defined as the mean hand angle during the 10 trials of the baseline block, movements that had been performed with congruent feedback (1^st^ target: trials 1 – 10; 2^nd^ target: trials 166 – 175). We defined three measures of learning: Early adaptation, late adaptation, and aftereffect. Early adaptation was operationalized as the mean hand angle over the first 10 trials of the perturbation block (1^st^ target: trials 11 – 20; 2^nd^ target: 176 – 185). Late adaptation was defined as the mean hand angle over the last 10 trials of the perturbation block (1^st^ target: trials: 151 – 160; 2^nd^ target: 316 – 325). The aftereffect was operationalized as the mean hand angle over the five trials of the no-feedback aftereffect block (1^st^ target: trials 161 – 165; 2^nd^ target: trials 326 – 330).

For each dependent variable, the data were submitted to a linear mixed effect model, with Rotation Size (3°, 10°, 30°, 45°) and Group (PD or control) as fixed effects and Participant ID as a random effect. As in Experiment 1, Years of Education was included as a covariate.

### Attention and Instruction Checks

It is difficult to verify that participants fully attend to the task in online studies. To address this issue, we sporadically instructed participants to make a specific keypress by presenting the message, “Press the letter “b” to proceed.” If the participant failed the make the instructed keypress, the experiment was terminated. These attention checks were randomly introduced within the first 50 trials of the experiment. We also wanted to verify that the participant understood the non-contingent feedback manipulation. To this end, we included the following instruction check after the three demonstration trials: “Identify the correct statement. Press ‘a’: I will aim away from the target and ignore the white dot. Press ‘b’: I will aim directly towards the target location and ignore the white dot.” The experiment was terminated if participants failed to make the correct response (i.e., press “b”). None of the participants failed these checks.

### Data Analysis

Outlier responses were defined as trials in which 1) the hand angle deviated by more than three standard deviations from the mean change in hand angle for each rotation size (Experiment 1) or from the moving average using a five-trial window, 2) the hand angle was greater than 75° from the target or 3) movement time exceeded 500 ms. These outlier trials were excluded from further analysis since behavior on these trials could reflect attentional lapses, anticipatory movements to another target location, or online corrections midway through the movement. The average percent of outlier trials were 7.6% ± 6.3% in Experiment 1 and 1.6 ± 2.2% in Experiment 2. This percentage is higher than in typical in-person studies, a pattern that has been observed previously in online studies with young adults (26).

In both experiments, we asked whether the learning measures for the PD group were correlated with motor symptom severity (UPDRS) and/or cognitive status (MoCA) (R function: cor.test). These variables were not included as covariates in the linear mixed effect model since they were only assessed in the PD group.

We employed F-tests with the Satterthwaite method to evaluate whether the coefficients (i.e., beta values) obtained from the linear mixed effects model were significant (R function: anova). Pairwise post-hoc t-tests (two-tailed) were used to compare the learning measures between the PD and Control groups (R function: emmeans). P-values were adjusted for multiple comparisons using the Tukey method and 95% confidence intervals are reported in squared brackets. Standard effect size measures were also provided (*D* for between-participant comparisons; *D_z_* for within-participant comparisons; 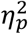 for between-subjects ANOVA) (40). We evaluated key null results using a Bayes Factor, BF_01_.

### Meta-analysis of the Effect of PD on Visuomotor Adaptation

We performed a meta-analysis of the effect of PD on visuomotor adaptation. For this analysis, we used searches through Google Scholar using the keywords: “Parkinson’s Disease,” “visuomotor adaptation”, “visuomotor rotation,’’ and “prism adaptation”. Study titles and abstracts were examined by three independent researchers (JT, LS, TN) who applied the following inclusion criteria: 1) The study had to employ a visuomotor adaptation task (e.g., prism adaptation, visuomotor rotation); 2) To control for medication state, the PD participants needed to be tested on their normal medication regimen (this led to exclusion of three experiments in (18,41,42); 3) The study had to include a dependent variable based on data from a post-perturbation phase (aftereffect). We used Cohen’s D as our measure of effect size for between-group comparisons (PD vs Controls). We opted to include the third criterion to focus on a measure of implicit adaptation less “contaminated” by strategy use.

The search yielded 4320 results from Google Scholar. Ultimately, 12 studies reporting the results from 16 experiments (253 total participants) met our three inclusion criteria. Of these 16 experiments, three were unpublished (also known as “gray literature”). The inclusion of unpublished work in a meta-analysis is recommended as one way to reduce publication bias (43).

For each of the 16 experiments, we calculated the aftereffect in one of three ways. 1) When we had access to the data, we calculated the effect size directly using the no-feedback washout trials (McDougle, Butcher, and Taylor’s unpublished observations, Exp 1 in this manuscript); 2) When included in the publication, we used the reported effect size (17,41,42,44,45). 3) When the effect size could not be directly inferred from the reported statistics but the washout data were presented graphically, we used Webplot Digitizer (https://automeris.io/WebPlotDigitizer/) to extract the aftereffect from the relevant figure, and used those values to calculate the effect size (11,18,19,46–48).

These aftereffect measures may not always provide a clean measure of implicit adaptation for two reasons. First, the magnitude of the aftereffect may depend on the instructions, and in particular whether participants are instructed to terminate the use of an re-aiming strategy and reach directly towards the visual target (49). If this instruction is not specified, aftereffects may measure both implicit adaptation and residual strategy use (50,51). Second, the time course of the aftereffect will depend on whether (veridical) visual feedback is provided. When provided, participants may become aware of their adapted state, and this could elicit a re-aiming strategy back towards the target. To focus on studies providing the “purest” measure of implicit adaptation, we performed a secondary analysis with a stricter inclusion criterion. Here we only included studies in which the participants were instructed at the start of the aftereffect block to stop using a strategy and reach directly to the target.

## Results

### Experiment 1: The Impact of PD on Implicit Adaptation

We asked how Parkinson’s Disease (PD) impacts implicit adaptation in response to a wide range of error sizes. To address this question, we varied the perturbation direction and the size of the non-contingent visual feedback using a random perturbation schedule throughout the perturbation block (Figure 1a). We assayed implicit adaptation by measuring the change in hand angle from trial n to trial n + 1 as a function of the rotation size on trial n. Positive values indicate a change in hand angle opposite to the direction of the visual feedback (i.e., an implicit adaptive response).

As can be seen in Figure 1b, the PD (green) and Control (dark magenta) participants exhibited an adaptive response, with the mean trial-to-trial change in hand angle in the opposite direction of the perturbation. This pattern was observed for all rotation sizes. Statistically, we first confirmed that the change in hand angle for each rotation size was significantly different than zero (all *t*_35_ > 4.9, *p* < 0.01; *D* > 0.8; 3°: 0.1, [0.6, 1.6]; 10°: 0.9, [1.5, 2.4]; 30°: 3.1, [2.3, 3.9]; 45°: 3.3, [2.7, 3.9]). Moreover, the magnitude of these trial-to-trial motor corrections converged with those observed in previous in-lab (35,52,53) and online studies (26) of implicit adaptation

Implicit trial-to-trial motor corrections are known to increase with perturbation size only within a limited range, saturating in response to larger perturbations (26,34,54–60). This sublinear “motor correction” function is thought to reflect an upper bound to trial-by-trial plasticity in either the sensory (61) or motor system (62). Visual inspection of the data (Figure 1b) indicated that the shape of the motor correction function was sublinear, increasing from 0° - 30° but saturating between 30° - 45°. Statistically, we verified this phenomenon in two ways: First, there was main effect of Rotation Size (*F*_1,106_ = 48.8, *p* < 0.001, *β* = 0.05, [0.03, 0.07], 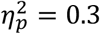), with the magnitude of the adaptive response increasing with the size of the perturbation. Second, the slope values computed with all rotation sizes was smaller than the slope values computed only with rotation sizes in the visually-defined linear zone (3°, 10°, 30°), indicating that the motor correction functions were sublinear (*t*_35_ = 2.3, *p* = 0.03, *β* = 0.02, [0.0, 0.04], *D_z_* = 0.4).

Turning to our main question, we next asked whether implicit adaptation would be impacted by PD. The main effect of Group (*F*_1,69.2_ = 0.0, *p* = 0.89, *β* = −0.01, [−1.2, 1.0], 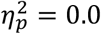) and the Group x Rotation Size (*F*_1,106_ = 0.0, *p* = 0.93, *β* = 0.0, [−0.03,0.03], 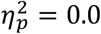) were not significant. A *post-hoc* comparison of the slopes showed a similar null effect, with a Bayes factor providing evidence in favor of the null hypothesis (*t*_34_ = −0.1, *p* = 0.93, [−0.04, 0.03], *D* = 0.0, BF_01_ = 3.1). In summary, the results of Experiment 1 indicate that implicit adaptation is preserved in PD across a wide range of rotation sizes.

### Experiment 2: The Impact of PD on the Upper Bound of Implicit Adaptation

Experiment 2 provided a second test of the effect of PD on implicit adaptation. To measure the time course of learning, we used clamped non-contingent feedback with the size of the clamp set to 3° or 30° in separate phases of the experiment (Figures 2a,d). In this manner, we obtain a picture of the cumulative effects of learning, an approach that should magnify any subtle differences between groups.

During the perturbation block, there was a gradual change in hand angle in the opposite direction of the clamped feedback, with the group-averaged functions approaching an asymptotic level after 50-100 clamped trials (Figure 2b,e). The adapted response was largely maintained during the following no-feedback block, consistent with what would be expected if learning induced by the clamped feedback is implicit and automatic.

Both groups exhibited robust changes in hand angle in all three phases of the experiment (t-test against 0: Early adaptation, PD: *t*_31_ = 6.7, *p* < 0.001, [6.8,12.7]; Controls: *t*_31_ = 5.8, *p* < 0.001, [5.9,12.3]; Late adaptation, PD: *t*_31_ = 7.3, *p* < 0.001, [20.0,35.4]; Controls: *t*_31_ = 9.8, *p* < 0.001, [15.4,23.4]; Aftereffect, PD: *t*_31_ = 7.3, *p* < 0.001, [15.3,27.2]; Controls: *t*_31_ = 8.4, *p* < 0.001, [11.9,19.7]). Implicit adaptation scaled with the size of the rotation during early adaptation (main effect of Rotation Size: *F*_1,30_ = 20.1, *p* < 0.001, *β* = 0.3, [0.1, 0.5], 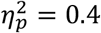), but reached similar values for the 3° and 30° clamps in late adaptation (F_1,30_ = 0.7, *p* = 0.51, *β* = 0.1, [−0.2,0.4], 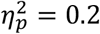) and in the aftereffect phase (*F*_1,30_ = 1.0, *p* = 0.33, *β* = 0.1, [−0.1,0.4], 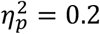). This asymptotic convergence is similar to that reported in Kim et al. (62) (but see: (51,63))

The learning functions for the PD and Control groups were statistically indistinguishable across all phases of the experiment, a null pattern that was observed in response to both the 3° and 30° clamps (Figures 2c,f). The main effect of Group and the Group x Rotation Size interaction were not significant for all three dependent variables (main effect of Group, Early: *F*_1,58_ = 0.1 *p* = 0.76, *β* = 1.0, [−4.9,6.9], 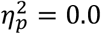; Late: *F*_1,53_ = 0.2, *p* = 0.70, *β* = 2.5, [−9.9,14.9], 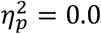; Aftereffect: *F*_1,50_ = 0.4, *p* = 0.51, *β* = 3.6, [−6.6,13.7], 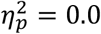; Group x Rotation Size interaction, Early: *F*_1,30_ = 0.1, *p* = 0.78, *β* = 0.0, [−0.3,0.2], 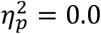, BF_01_ = 1.0; Late: *F*_1,30_ = 1.9, *p* = 0.07, *β* = 0.4, [0.0,0.9], 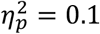, BF_01_ = 0.5; Aftereffect: *F*_1,30_ = 1.8, *p* = 0.19, *β* = 0.2, [−0.1,0.6], 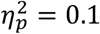, BF_01_ = 0.3). If anything, there was a trend towards greater implicit adaptation in the PD group, driven by a few “super adapters” (i.e., individuals with a heading angle that deviated by more than 25° from the target location during late adaptation). Nonetheless, our meta-analysis of the literature (see next section) did not provide any evidence in favor of this trend.

In summary, the results of Experiment 2 converge with those obtained in Experiment 1, providing additional evidence that implicit adaptation to a visuomotor perturbation is preserved in PD for both small and large error sizes.

### Meta-analysis of the Effect of PD on Visuomotor Adaptation

We conducted a meta-analysis of previous studies examining the effect of PD on visuomotor adaptation. Most of these studies used a standard perturbation, one in which the position of the feedback is contingent on the participant’s hand position. As such, adaptation results in improved performance with the cursor landing closer to the target. Recent work has shown that performance in such tasks is influenced, and even dominated (for large perturbations) by explicit re-aiming strategies rather than implicit adaptation (20,21,44,50,64–68). As such, we focused on studies that included an aftereffect phase in which the perturbation was removed.

A summary of the meta-analysis is presented in Figure 3. Positive effect sizes indicate results in which the aftereffect measure was larger for the Control group compared to the PD group (i.e., impaired adaptation in PD); negative values indicate a greater aftereffect for the PD group. When the confidence interval for a given study includes 0, the group comparison was not significant. This null pattern holds in 12 of the 16 experiments, and the grand effect size also encompasses 0 (*D* = 0.33 [−0.05,0.71]). Thus, the overall pattern in the meta-analysis indicates that implicit adaptation is not affected by PD.

**Figure 3:**
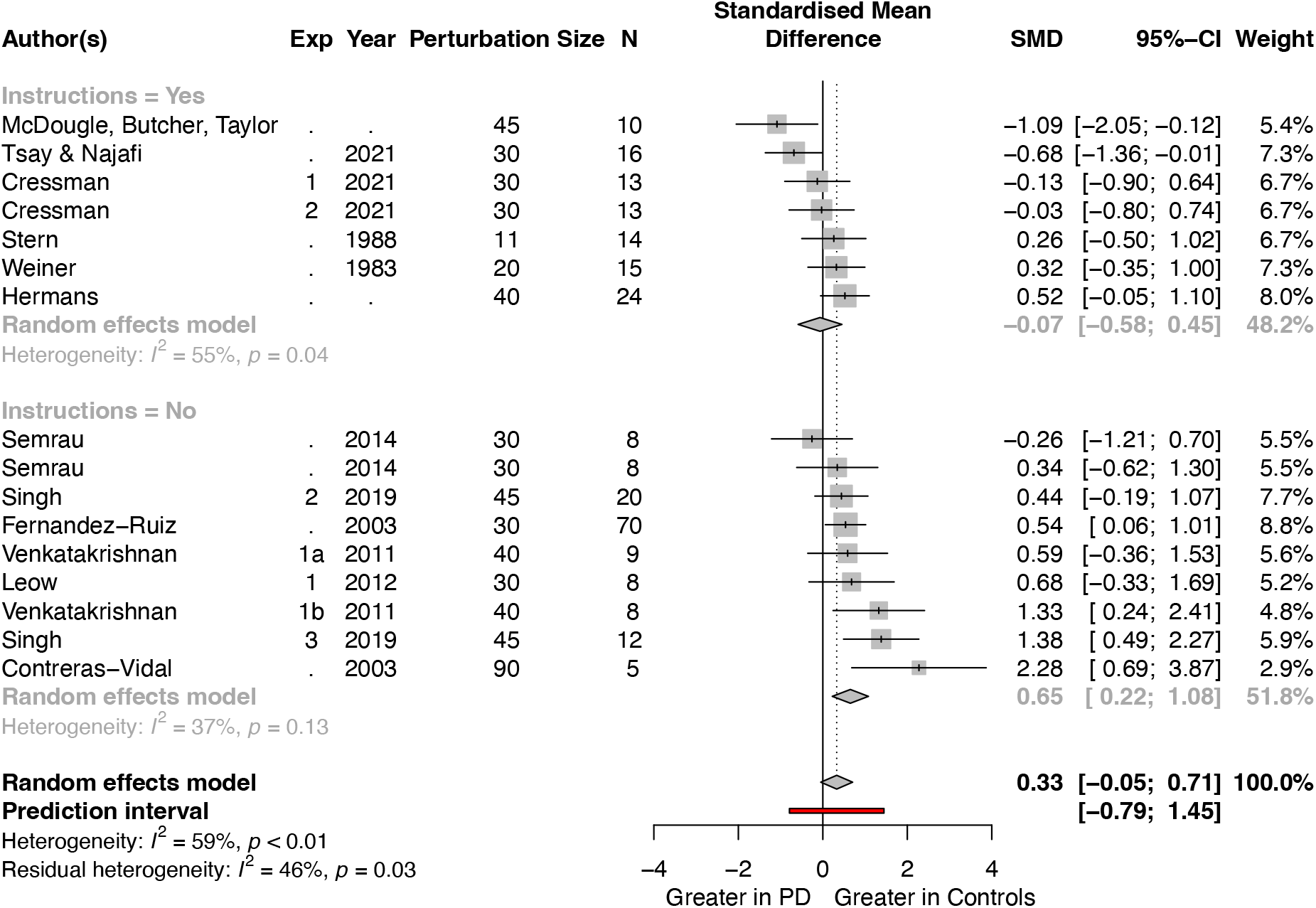
Implicit adaptation is preserved in PD over a wide range of studies. The data for this meta-analysis are taken from 16 experiments reported in 12 papers. Studies are organized according to whether the instructions at the start of the washout block explicitly emphasized that participants should stop using a strategy (top) or were ambiguous (bottom). All participants were tested on their regular medical schedule and (when applicable) with DBS turned on. The grand effect size, along with the 95% CI, across all of the experiments is indicated at the bottom of this forest plot (diamonds).

Note that we opted to only include studies examining PD participants on their normal medication regimen to 1) control for medication schedule and 2) avoid double dipping, that is, counting the same individual twice – on and off medication– in the same meta-analysis (see: (18,41,42)). Nonetheless, there were three visuomotor adaptation studies in which aftereffect data were available for PD participants who had been tested off medication. The experiments reported in Singh et al (2019) found attenuated aftereffects in PD tested off medication (*D* = 2.61 [1.77,3.46]) or off DBS (*D* = 2.20 [0.27,3.22]). However, the results from the other two studies indicate null effects (Cressman et al (2021), Exp 1: *D* = −0.48 [−1.26,0.30]; Exp 2: *D* = 0.23 [−0.53,1.01]; Semrau et al (2014), Clockwise rotation, off-med: *D* = −1.49 [−2.57, −0.41]; Counterclockwise rotation, off-med: *D* = 0.30 [−0.66,1.26]). Although the sample size here is quite limited, these results suggest that implicit adaptation remains intact in individuals with PD even when tested outside their normal medical regimen.

We recognize that aftereffect measures may not always provide a clean measure of implicit adaptation. For example, if participants are not explicitly told to reach directly to the target, terminating the use of any strategy they may have adopted, aftereffect performance may measure both implicit adaptation and residual strategy use (48,50,51). The significant heterogeneity (*I*^2^ = 59%, *p* < 0.01) in the 16 experiments may, in part, be driven by differences in the instructions provided prior to the washout block. To focus on studies providing the “purest” measure of implicit adaptation, we added a stricter inclusion criterion in a secondary analysis, requiring that the participants had been instructed to stop using a strategy and reach directly to the target during the aftereffect trials. Seven experiments remained after we imposed this additional criterion (top half of Figure 3). Six of the seven studies observed no group difference, and the other showed greater implicit adaptation in PD. The grand effect size comparing the PD and Control groups again encompassed 0 (*D* = −0.07 [−0.58,0.45]). Together, this subgroup analysis implies that implicit adaptation, when cleanly isolated, is not impaired in PD.

The studies that did not clearly instruct participants to stop using their strategy are shown in the bottom portion of Figure 3. Here, there was a group difference (*D* = 0.65 [0.22,1.08]), with the control participants exhibiting a larger aftereffect than the PD participants. Notably, the PD participants in most of these studies also showed attenuated performance during the trials right before the aftereffect block (i.e., late adaptation), a phase in which the use of an explicit re-aiming strategy is encouraged. As such, a parsimonious interpretation of these data is that the late adaptation deficit is due to impairment in strategic aiming, and the corresponding aftereffect deficit is also due to greater residual strategy use in the control participants.

Taken together, the meta-analysis results from the instruction-based subgroups points to a dissociation whereby PD does not impact implicit adaptation but does impair more explicit aspects of performance on visuomotor adaptation tasks.

## Discussion

The basal ganglia are an integral part of motor system, contributing to the acquisition and automatization of skilled movements. Studies involving individuals with Parkinson’s Disease have reinforced this notion, showing deficits in a wide range of motor learning tasks (4–10). Here, we homed in on a critical component of sensorimotor learning, evaluating the integrity of implicit adaptation in individuals with PD. Although this topic has been addressed in many studies, an assortment of methodological issues has precluded a clear answer on the basic question of whether implicit adaptation is disrupted in PD.

In revisiting this question, we used non-contingent visual feedback in a visuomotor adaptation task, a method in which performance changes are completely implicit; as such, this method isolates implicit motor adaptation without the influence of cognitive strategies (23). Using both variable (Exp 1) and fixed (Exp 2) perturbations, we found that the form and extent of implicit adaptation was indistinguishable between PD and controls. This null effect was observed in response to a wide range of error sizes (3° - 45°). We complemented our experiments with a meta-analysis, identifying 16 experiments involving over 200 PD participants that included a relatively clean measure of implicit adaptation (i.e., aftereffect). The overall pattern in these studies, as well as the aggregated effect size, indicated that implicit adaptation was not affected by PD.

We recognize that the literature does include positive results, cases in which PD participants were impaired on sensorimotor adaptation tasks. Some of the positive results could reflect Type I errors, a problem that is amplified considering the substantial within-subject variability as well as relatively small sample sizes typical in most neuropsychological studies. However, it is also important to consider if the observed deficits attributed to implicit motor adaptation actually reflect impairment in the utilization of explicit strategic processes (20,21). While we tried to minimize this possibility by focusing our meta-analysis on performance during the washout phase when the perturbation has been removed, aftereffect measures may still be contaminated if the participants continue to apply an aiming strategy. This concern is especially relevant if the instructions fail to emphasize that the participant should stop aiming and reach directly to the target (49). As can be seen in Figure 3, this concern may be relevant for over half of the studies included in our meta-analysis. If the meta-analysis is restricted to studies in which participants were explicitly instructed in the washout phase to reach directly to the target (i.e., terminate the use of an aiming strategy), we observe a null effect, providing converging evidence that implicit adaptation is preserved in PD.

In contrast, the meta-analysis suggests that participants with PD may be impaired in deriving and/or applying an aiming strategy to counteract a perturbation. Control participants exhibited a larger aftereffect when the instructions failed to instruct participants to stop using an aiming strategy. Under such conditions, explicit aiming likely contributes to the aftereffect. A finer-grain analysis of the two studies that show the largest PD deficit supports this hypothesis. Contreras et al (17) found that, following exposure to a 90° rotation, the PD group exhibited a marked reduction in the magnitude of the aftereffect relative to a control group. However, the aftereffect for the control group is likely contaminated by explicit strategies: The mean was three times larger (~60°) than that typically observed when participants are instructed to reach directly to the target during washout (~20°) (62,69,70). Moreover, a similar degree of impairment in the PD group was observed during late adaptation, a phase in which the use of an explicit strategy to counteract the perturbation is essential for successful performance. A similar pattern is found in Singh et al (2020) where the perturbation was a 45° rotation: Both late adaptation and aftereffects show a similar impairment in PD, with the control group again showing an approximate two-fold increase in the aftereffect size compared to the typical value found in the literature when steps are taken to eliminate explicit contributions to the aftereffect. Thus, the difference between the PD and control groups may reflect PD-related impairment in strategic aiming, with the control group using a more effective aiming strategy than the PD group, rather than a PD-related deficit in implicit adaptation.

Consideration of other phenomena observed in studies of sensorimotor adaptation also point to an impairment in explicit strategy use in PD. For instance, savings, a phenomenon characterized by accelerated learning upon re-exposure to a perturbation after washout or a long break (16,65,71–73), is impaired in PD (74) (but see: (46)). Recent studies have shown that savings arises from the faster recall of a previously learned strategy (71); in contrast, savings is not observed in measures of implicit adaptation (75). A related phenomenon involves tasks in which the participant must successively learn multiple visuomotor mappings (e.g., 45° clockwise rotation block followed by 45° counterclockwise rotation block). Performance on these tasks is facilitated by the successful utilization of multiple aiming strategies (76). Here, too, an impairment has been observed in PD (77). Moreover, learning large perturbations (e.g., 90° rotations) that are suddenly introduced have been shown to demand greater strategy use than learning small perturbations that are gradually introduced (78,79). In one study, PD participants were found to be impaired in the former, but not in the latter (19). Future studies using methods that directly probe explicit re-aiming (see: (20,50)) can directly test strategy use in PD.

The absence of a PD-related impairment in implicit sensorimotor adaptation stands in contrast to the marked impairment observed in individuals with spinocerebellar degeneration on this form of learning (80,81). This cerebellar-related deficit has been observed across a wide range of tasks using different perturbation sizes (82), perturbation schedules (83), perturbation types (80,84), and effectors (85–87). For example, using clamped feedback, Morehead et al (2017) found that implicit adaptation was attenuated by ~50% in a group of individuals with cerebellar degeneration. Taken together, there is a clear dissociation between the effects of degenerative processes impacting the basal ganglia or cerebellum: Implicit adaptation appears to depend on the integrity of the cerebellum and not the basal ganglia.

It remains to be seen if the insights gleaned from the current study also call into question other lines of evidence indicating a PD-related impairment in implicit sensorimotor learning. Our empirical findings and the meta-analysis highlight the challenge faced when using tasks as models of specific learning processes: Namely, that these tasks likely involve multiple learning processes and care must be taken to isolate the contribution of each process as well as the interaction between different processes (see also: (88)). To take one example, studies of the effect of PD on sequence learning have also yielded ambiguous results: A number of studies have reported deficits in sequence learning in PD (4,7,27), while other have reported no impairment (89–91). However, it has long been recognized that measures of sequence learning frequently involve the combined effects of implicit and explicit processes (92,93). Similar to the approach taken here, it would be important to revisit sequence learning with methods or tasks that provide purer measures of implicit sequence learning (e.g., see (4,94,95)). Even the acquisition of simple stimulus-response contingencies and mirror drawing do not rely exclusively on implicit processes to support incremental error-based learning, but can involve higher level functions that derive heuristics to facilitate learning (96–101). It will be interesting to explore if the impaired performance observed in PD participants on tasks traditionally thought to reflect implicit learning may instead reflect an impairment in other, more explicit forms of learning.

## Acknowledgements

We thank Sam McDougle, Jordan Taylor, and Lucio Marinelli for sharing their data for our meta-analysis. We also thank Sharon Binoy, Rachel Woody, and Will Saban for conducting neuropsychological assessments via the PONT platform (27)

## Data availability statement

Data will be available upon publication at https://github.com/xiaotsay2015.

## Supplementary Covariate Analyses

### Experiment 1, covariate analysis of the PD group

The degree of implicit adaptation was not associated with years of education (*F*_1,33_ = 0.4, *p* = 0.53, *β* = −0.05, [−0.2,0.1], 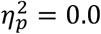), MoCA score (slope computed with all rotation sizes vs MoCA: *R* = 0.1, *p* = 0.66, [−0.4,0.6]), or UPDRS score (slope with all errors vs UPDRS: *R* = 0.1, *p* = 0.79, [−0.4, 0.5]).

### Experiment 2, covariate analysis of the PD group

Implicit adaptation was not associated with years of education (early: *F*_1,29_ = 0.1, *p* = 0.74, *β* = 0.1, [−0.6, 0.8], 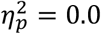; late: <u>*F*_1,29_ = 0.6, *p* = 0.43, *β* = −0.7, [−2.3.1.0]</u>, 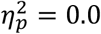; aftereffect: : *F*_1,29_ = 1.7, *p* = 0.20, *β* = −0.9, [−2.3,0.5]), MoCA score (early: *R* = −0.1, *p* = 1, [−0.4,0.3]; late: *R* = 0, *p* = 1, [−0.3, 0.4]; aftereffect: *R* = 0.39, *p* = 0.46, [−0.2, 0.5]), or UPDRS score (early: *R* = −0.1, *p* = 1, [−0.4, 0.3], late: *R* = −0.2, *p* = 0.75, [−0.5,0.2]; aftereffect: *R* = −0.39, *p* = 0.12, [−0.6,0.0]).

| Experiment 1 |  |  |  |  |  |  |  |  |  |  |
| --- | --- | --- | --- | --- | --- | --- | --- | --- | --- | --- |
| Parkinson's Disease (n = 18) |  |  |  |  |  |  | Control (n = 18) |  |  |  |
| E1_ID | Age | Sex | Hand | MoCA | Education | UPDRS | Age | Sex | Hand | Education |
| 1<sup>#</sup> | 72 | M | R | 27 | 21 | 16 | 71 | M | R | 16 |
| 2<sup>#</sup> | 69 | M | R | 30 | 16 | 12 | 70 | M | R | 16 |
| 3<sup>#</sup> | 81 | M | R | 25 | 27 | 21 | 78 | M | R | 18 |
| 4<sup>#</sup> | 58 | F | L | 26 | 18 | 9 | 58 | F | L | 16 |
| 5<sup>#</sup> | 70 | F | R | 28 | 18 | 11 | 67 | F | R | 16 |
| 6 | 65 | M | R | 30 | 16 | 21 | 65 | M | R | 20 |
| 7<sup>#</sup> | 52 | F | R | 28 | 16 | 21 | 54 | F | R | 18 |
| 8 | 67 | F | R | 30 | 18 | 16 | 64 | F | R | 12 |
| 9<sup>#</sup> | 64 | M | R | 27 | 20 | 25 | 64 | M | R | 16 |
| 10 | 58 | F | R | 30 | 18 | 20 | 56 | F | R | 12 |
| 11 | 76 | M | R | 30 | 18 | 21 | 74 | M | R | 16 |
| 12 | 53 | M | L | 29 | 12 | 21 | 52 | M | L | 16 |
| 13 | 63 | F | A | 25 | 14 | 12 | 62 | F | A | 13 |
| 14 | 73 | F | R | 28 | 19 | 10 | 69 | F | R | 14 |
| 15 | 66 | F | R | 28 | 16 | 16 | 64 | F | R | 12 |
| 16<sup>#</sup> | 64 | M | R | 27 | 14 | 20 | 61 | M | R | 14 |
| 17<sup>#</sup> | 70 | F | L | 30 | 16 | 10 | 67 | F | L | 14 |
| 18 | 61 | F | R | 30 | 16 | - | 58 | F | R | 16 |
| Mean (SD) | 65.7 (7.6) | 8 M<br>10 F | 14 R<br>3L, 1 A | 27.8 (1.8) | 17.4 (3.3) | 16.6 (5.1) | 64.1 (6.9) | 8 M<br>10 F | 14 R<br>3L, 1 A | 15.3 (2.2) |
| Experiment 2 |  |  |  |  |  |  |  |  |  |  |
| Parkinson's Disease (n = 16) |  |  |  |  |  |  | Control (n = 16) |  |  |  |
| E2_ID | Age | Sex | Hand | MoCA | Education | UPDRS | Age | Sex | Hand | Education |
| 1<sup>#</sup> | 71 | M | R | 27 | 21 | 16 | 72 | M | R | 16 |
| 2<sup>#</sup> | 63 | M | R | 27 | 14 | 20 | 64 | M | R | 13 |
| 3 | 59 | M | R | 30 | 18 | 10 | 60 | M | R | 14 |
| 4<sup>#</sup> | 69 | F | L | 30 | 16 | 10 | 64 | F | L | 16 |
| 5 | 66 | M | R | 25 | 18 | 21 | 62 | M | R | 16 |
| 6<sup>#</sup> | 68 | M | R | 30 | 16 | 12 | 68 | M | R | 16 |
| 7<sup>#</sup> | 80 | M | R | 25 | 27 | 21 | 78 | M | R | 18 |
| 8<sup>#</sup> | 57 | F | L | 26 | 18 | 9 | 57 | F | L | 16 |
| 9<sup>#</sup> | 69 | F | R | 28 | 18 | 11 | 68 | F | R | 14 |
| 10<sup>#</sup> | 64 | M | R | 30 | 16 | 21 | 61 | M | R | 14 |
| 11 | 68 | F | R | 25 | 16 | 15 | 68 | F | R | 16 |
| 12<sup>#</sup> | 51 | F | R | 28 | 16 | 21 | 48 | F | R | 16 |
| 13 | 73 | F | R | 28 | 20 | 25 | 73 | F | R | 12 |
| 14 | 77 | F | L | 29 | 13 | 18 | 70 | F | L | 23 |
| 15 | 69 | F | R | 29 | 16 | 21 | 72 | F | R | 14 |
| 16 | 66 | F | R | 30 | 18 | 16 | 63 | F | R | 12 |
| Mean (SD) | 66.9 (7.2) | 7 M<br>9 F | 13 R<br>3 L | 27.4 (1.9) | 17.6 (3.2) | 16.7 (5.0) | 65.6 (7.2) | 7 M<br>9 F | 13 R<br>3 L | 15.4 (2.6) |

